# Comparative genomics of clinical isolates of *Pseudomonas aeruginosa* from cystic fibrosis patients in Mexico

**DOI:** 10.64898/2026.08.28.747926

**Authors:** Evelin Martínez-Rosales, Armando Gerónimo-Gallegos, Francisco Cuevas Schacht, Miriam Sarahi Lozano Gamboa, Marisol López-López, Rodolfo García-Contreras, Rafael Coria-Jiménez, Corina Diana Ceapă

## Abstract

*Pseudomonas aeruginosa* (*P. aeruginosa*) is the primary pathogen responsible for morbidity and mortality in patients with cystic fibrosis (CF). Its genomic plasticity and constant selective pressure from antimicrobial treatments have favored the emergence of multidrug-resistant clones. This study conducted a comparative genomic analysis of 41 *P. aeruginosa* isolated from pediatric patients with CF in Mexico from 2015 to 2024, with the aim of characterizing their evolutionary dynamics, resistome, and virulome.

Whole-genome sequencing (MGI, Illumina, and PacBio platforms) was used, with *de novo* assemblies performed using Unicycler v0.4.8 on the BV-BRC platform. The databases used for the resistome were CARD and NDARO, and for the virulome, VFDB. Phylogenetic reconstruction was based on core-genome alignments generated with Roary v3.13.0, with maximum likelihood reconstruction performed in IQ-TREE v2.1.2. The statistical significance of the segregation of resistance and virulence patterns was evaluated using PERMANOVA analysis.

The results revealed a significant clonal prevalence of sequence types (ST) 307 and ST 167. Phylogenomic analysis grouped the isolates into three main clades; Clade 1 stood out for having the highest resistance gene load (mean of 75 genes/genome), establishing itself as the main reservoir of multidrug-resistant profiles. Genotype-phenotype concordance reached 65.5% overall, with high accuracy for aminoglycosides (87.8%) and fluoroquinolones (82.9%). Furthermore, virulome analysis identified 67 distinct patterns that were significantly segregated among the clades (PERMANOVA: *R^2^*=0.31, *p*=0.001). These findings demonstrate that the evolution of *P. aeruginosa* lineages in the pediatric clinical setting involves parallel and coordinated adaptations in both their resistance potential and their virulence arsenal.

This study underscores the need to adopt a multidisciplinary approach to the clinical management of chronic *P. aeruginosa* infections in pediatric patients. The persistence of extensively drug-resistant (XDR) strains calls for the integration of genomic surveillance and functional diagnostics, as well as the search for therapeutic alternatives for the clinical management of patients with cystic fibrosis.

**Impact statement:** This study identifies high-risk, circulating *P. aeruginosa* lineages in pediatric patients with cystic fibrosis in Mexico. By demonstrating the parallel evolution of the resistome and virulome, we provide a genomic framework that enhances our understanding of chronic infection persistence. These findings offer a critical foundation for optimizing the surveillance of circulating strains and advancing therapeutic strategies to improve patient outcomes.

**Data summary:** All genomic data generated in this study have been deposited in the National Center for Biotechnology Information (NCBI) database under the BioProject accession number PRJNA1439432. Specific accession numbers for each isolate are listed in Table S1 (Supplementary Data). The authors confirm that all supporting data, code, and protocols have been provided within the article or through supplementary data files.

**Repositories:** All genome assemblies and raw sequencing data have been deposited in the NCBI BioProject database under accession number PRJNA1439432. Individual isolate accession numbers are listed in Table S1.

## Introduction

*Pseudomonas aeruginosa* (*P. aeruginosa*) is a Gram-negative, facultatively aerobic bacterium and one of the leading causes of severe infections in immunocompromised individuals [1]. Its clinical adaptability is attributed to a predominantly circular genome ranging from 5.5 to 7 Mbp, with a guanine-cytosine content exceeding 65%, that encodes a broad repertoire of regulatory networks and metabolic pathways [2]. The close host-parasite association between *P. aeruginosa* and the patient with cystic fibrosis (CF) has been widely described. CF is a genetic disorder with multisystemic effects; however, its primary cause of morbidity and mortality stems from recurrent and chronic bacterial lung disease, which leads to progressive deterioration of lung function [1]. In this context, *P. aeruginosa* stands out as a highly virulent and antibiotic-resistant pathogen, possessing effective adaptation mechanisms that allow it to establish itself in the airways of patients with CF and develop chronic infection with high case-fatality rates [2].

*P. aeruginosa* represents a global health emergency due to the continuous emergence of multidrug-resistant strains, underscoring the limitations of traditional phenotypic diagnostic methods [3]. Consequently, genomic analysis has become essential for deciphering the underlying mechanisms of persistence and virulence, providing a critical framework for the development of targeted antimicrobial and anti-virulence therapeutic alternatives [4]. Whole-genome sequencing (WGS) of *P. aeruginosa* in the context of CF provides insight into the complete set of resistance and virulence genes—both intrinsic and accessory—in strains with versatile adaptive capabilities, addressing a clinical need to understand the trajectory of strains that establish chronic infections [5].

Given the clinical burden of chronic *P. aeruginosa* infection in CF and the limited genomic data available from Latin American cohorts, it is essential to characterize clonal diversity, antimicrobial resistance determinants, and evolutionary trajectories. High-risk circulating lineages in Mexico pose a significant challenge for clinical management, and understanding the interaction between the evolution of the resistome and the virulome is essential for effective epidemiological surveillance. Therefore, the objective of this study was to conduct a comprehensive genomic analysis of clinical *P. aeruginosa* strains isolated from pediatric patients in Mexico. We applied an integrated comparative genomics approach to characterize clonal diversity, antimicrobial resistance determinants, virulence repertoires, and evolutionary trajectories in isolates collected over nearly a decade. By integrating phylogenomic reconstruction with resistome and virulome profiling, we sought to elucidate the adaptive strategies of these pathogens, providing a robust genomic framework to improve clinical surveillance and guide the development of more effective and personalized therapeutic interventions.

## Methods

### Ethics and Biosafety

Respiratory samples were collected as part of routine clinical care and processed under institutional approval (INP-2026/011), following national regulations (NOM-012-SSA3-2012) and international guidelines for genomic data handling. All laboratory work involving *P. aeruginosa* was conducted under BSL-2 conditions using validated institutional SOPs.

### Bacterial isolates and clinical metadata

*P. aeruginosa* isolates were obtained from paediatric cystic fibrosis (CF) patients treated at the National Institute of Pediatrics (Mexico City) between 2015–2024 [6]. Samples included sputum, bronchoalveolar lavage (BAL), and nasopharyngeal aspirates. Species identification followed ASM recommendations [3]. Clinical metadata included patient age, sex, CFTR genotype (when available), sample type, year of isolation, and coinfection profile. Longitudinal isolates were available for several patients, enabling within-host evolutionary comparisons.

### Antibiotic susceptibility testing

Minimum inhibitory concentrations (MICs) were determined using Liofilchem® MIC Test Strips according to CLSI M100 (30th ed., 2024) [4]. Bacterial suspensions (1–2 × 10 CFU/mL) were inoculated onto Mueller–Hinton agar, and MICs were read after 18–24 h at 37 °C by interpreting the intersection of the inhibition ellipse with the strip scale.

### Selection of strains for sequencing

A total of 41 *P. aeruginosa* strains were selected for Whole Genome Sequencing (WGS). The selection criteria were the bacterial susceptibility and resistance pattern to antibiotics, the patient from whom the isolate was obtained, and the year of isolation, with priority given to highly resistant microorganisms and strains isolated from the same patient over the years.

### Genome sequencing and assembly

Genomic DNA from the 41 *P. aeruginosa* clinical isolates was extracted from overnight cultures grown in Luria Bertani media (LB) using standard phenol–chloroform-CTAB (cetyltrimethylammonium bromide) purification [7, 8]. DNA quantity and purity were assessed with Qubit fluorometry and NanoDrop spectrophotometry, and high-molecular-weight integrity was verified by agarose gel electrophoresis. Libraries were prepared according to the manufacturer’s protocols for each sequencing platform. Most isolates were sequenced using short-read technologies (MGI and Illumina), while a subset of representative strains underwent long-read sequencing with PacBio to improve assembly contiguity.

Raw reads were quality-filtered and adapter-trimmed before assembly. Datasets were assembled *de novo* using Unicycler version v0.4.8 [9] in the BV-BRC platform (Bacterial and Viral Bioinformatics Resource Center) [10]. Assembly quality was evaluated using contiguity metrics and genome completeness obtained with CheckM v1.2.2 [11]. Of the 51 sequenced isolates, 41 met the predefined quality thresholds, including fewer than 200 contigs, GC content between 65– 66%, estimated contamination below 10%, and total genome size between 6 and 7 Mb. These high-quality assemblies were retained for downstream comparative genomics analyses.

All genome assemblies and raw sequencing data have been deposited in the NCBI BioProject database under accession number PRJNA1439432 (Supplementary Table S1).

### Genome Annotation

To ensure uniformity across analyses, all genomes were re-annotated using Prokka v1.14.6 [12] (GFF3 output, genetic code 11, minimum contig length 200 bp) and RASTtk [13] (BV-BRC v3.30.19) for complementary functional annotation. Annotations were used as input for pangenome, phylogenomic, resistome, virulome, and mobilome analyses.

### Phylogenomic Analysis

Core-genome alignments were generated using Roary v3.13.0 [14] (95% BLASTP identity threshold; core genes ≥95% of genomes). Maximum-likelihood phylogenies were visualized with IQ-TREE v2.1.2 [15]. ModelFinder for best-fit substitution model (e.g., GTR+F+I+G4) and 1,000 ultrafast bootstrap replicates. Metadata (patient ID, year, sample type, MLST, sequencing platform) were integrated into tree visualizations using iTOL v6 [16].

### Multilocus Sequence Typing (MLST)

MLST profiles were assigned using the PubMLST *P. aeruginosa* scheme. Alleles for housekeeping genes were extracted using BLASTN and submitted to PubMLST for ST assignment. Novel allelic combinations were flagged and reported.

### Resistome Profiling

Resistome characterization followed the same analytical framework applied in our previous genomic surveillance studies. Genome assemblies were screened using the BV-BRC K-mer– based annotation pipeline and subsequently validated through BLAST searches against the CARD [17] and NDARO [18] repositories to ensure high-confidence detection of antimicrobial resistance determinants. To increase sensitivity and reduce false negatives, we incorporated the SraX workflow [19], configured with DIAMOND BlastX alignment [20], an identity threshold of 85%, and a minimum coverage of 60%. Outputs from all tools were consolidated into a non-redundant resistome matrix in which homologous genes were grouped, and conflicting annotations were resolved by prioritizing presence calls.

Resistome variation was quantified using Bray–Curtis dissimilarity indices derived from presence/absence matrices. Principal Coordinates Analysis (PCoA) and k-means clustering were used to identify resistome clusters; significance was assessed via PERMANOVA. Core resistance genes were defined as those present in ≥80% of genomes; rare determinants were those detected in ≤3 isolates.

### Virulome Profiling

Virulence genes were identified using BV-BRC annotations and curated virulence factor databases (VFDB) [21]. Presence/absence matrices were generated and compared across isolates. Virulome diversity was assessed independently of phylogeny to evaluate horizontal gene transfer and within-host adaptation.

### Mobile elements profiling

Mobile genetic elements were identified using **VRprofile2** [22], which enabled systematic detection of plasmids, insertion sequences, integrases, transposons, and other mobilization modules across all genomes. Each assembly was analyzed independently, and the resulting annotations were curated to quantify the abundance and diversity of MGE families. Mobilome composition was compared across isolates to identify patterns associated with longitudinal persistence, co-infection contexts, or specific phylogenetic backgrounds.

To explore the contribution of MGEs to genomic plasticity, we examined the co-localization of resistance and virulence determinants within mobile regions and assessed the distribution of replicon types and transposase families. Mobilome dynamics were visualized using SankeyMATIC to illustrate the flow of genetic material between chromosomes, plasmids, and mobile elements, providing an integrated view of horizontal gene transfer events within the cystic fibrosis airway environment.

### Statistical analyses

All statistical analyses were performed in R v4.3.2. PERMANOVA, clustering, and diversity metrics were computed using vegan and stats packages. Figures were generated using ggplot2, Cytoscape, and BV-BRC visualization tools.

## Results

### Clinical and epidemiological characterization of P. aeruginosa isolates from patients with CF

A total of 41 clinical isolates of *P. aeruginosa* from respiratory secretions of 16 patients with CF who attend pediatric appointments at the National Institute of Pediatrics in Mexico, collected from 2015–2024, were analyzed. The patients ranged in age from 7 months to 17 years, with an equal distribution of sexes (8 males and 8 females); although sputum was the most prevalent isolation source, bronchoalveolar lavage or nasopharyngeal suction were preferentially obtained when patients were in critical clinical conditions or unable to produce sputum, particularly in very young paediatric cases (Table 1).

**Table 1:** Clinical characteristics and MLST of *Pseudomonas aeruginosa* clinical isolates.

| Patients | Sex | CFTR mutation | Genome Name | Isolation year | Age | Isolation source | MLST |
| --- | --- | --- | --- | --- | --- | --- | --- |
| P1 | Male | Undetermined | Pa127r | 2020 | 11 | Sputum | 167 |
|  |  |  | Pa128m | 2021 | 12 | Sputum | 167 |
|  |  |  | Pa128r | 2021 | 12 | Sputum | 167 |
|  |  |  | Pa130m | 2021 | 12 | Sputum | 167 |
|  |  |  | Pa130mch | 2021 | 12 | Sputum | 167 |
|  |  |  | Pa131m | 2021 | 12 | Sputum | 167 |
| P2 | Female | p.Phe508del | Pa129m | 2021 | 15 | Sputum | 381 |
|  |  |  | Pa129r | 2021 | 15 | Sputum | 381 |
|  |  |  | Pa132m1 | 2021 | 16 | Sputum | 167 |
|  |  |  | Pa132m2 | 2021 | 16 | Sputum | 167 |
|  |  |  | Pa132r | 2021 | 16 | Sputum | 395 |
|  |  |  | Pa133r | 2022 | 16 | Sputum | 381 |
| P3 | Male | DI507 | Pa124ra | 2019 | 11 | BL<sup>1</sup> | 460 |
|  |  |  | Pa124rb | 2019 | 11 | BL | 309 |
| P4 | Male | Undetermined | Pa147r | 2023 | 13 | Sputum | 234 |
| P5 | Male | Undetermined | Pa152r | 2023 | 17 | Sputum | 644 |
|  |  |  | Pa162r | 2023 | 17 | BL | 644 |
| P6 | Male | Unknown | Pa112r | 2018 | 4 | BL | NT |
| P7 | Female | p.Phe508del,<br>7148T | Pa136m | 2023 | 13 | ND<sup>2</sup> | NT |
| P8 | Female | Unknown | Pa140r1 | 2023 | 9 m | NPA<sup>3</sup> | 395 |
|  |  |  | Pa140r2 | 2023 | 9 m | NPA | 395 |
| P9 | Female | C.1646G>A<br>heterozygous<br>status | Pa197m1 | 2024 | 9 | BL | NT |
| P10 | Male | NI303K | Pa122r | 2019 | 8 | Sputum | 1337 |
|  |  |  | Pa22r | 2015 | 5 | Sputum | 1565 |
| P11 | Male | Undetermined | Pa118ma | 2018 | 10 | Sputum | 235 |
|  |  |  | Pa169m | 2023 | 13 | Sputum | NT |
|  |  |  | Pa188m | 2024 | 14 | Sputum | NT |
| P12 | Female | I48T | Pa78r | 2017 | 13 | Sputum | 1565 |
| P13 | Female | Compound<br>heterozygote<br>G542X/X | Pa176r | 2024 | 14 | Sputum | 309 |
|  |  |  | Pa182r1 | 2024 | 14 | Sputum | 309 |
|  |  |  | Pa186m | 2024 | 15 | Sputum | 309 |
|  |  |  | Pa186r | 2024 | 15 | Sputum | 309 |
|  |  |  | Pa189m | 2024 | 15 | Sputum | 309 |
|  |  |  | Pa194m | 2024 | 15 | Sputum | 309 |
|  |  |  | Pa194r | 2024 | 15 | Sputum | 309 |
|  |  |  | Pa196r | 2024 | 15 | Sputum | 309 |
|  |  |  | Pa201m | 2024 | 15 | Sputum | 309 |
| P14 | Male | G542X | Pa70m1 | 2016 | 12 | BL | 309 |
|  |  |  | Pa83r | 2017 | 12 | Sputum | 309 |
| P15 | Female | Undetermined | Pa198r1 | 2024 | 7m | BL | 309 |
| P16 | Female | Unknown | Pa47r | 2016 | 7 | NA | 309 |

<sup>1</sup>Bronchoalveolar lavage, <sup>2</sup>No Data, <sup>3</sup>Nasopharyngeal aspirate

### Genomic epidemiology reveals a high clonal predominance of ST309 and ST167

Phylogenomic and MLST profiling revealed substantial clonal diversity among the isolates, identifying a total of 10 STs. High predominance of ST309 was observed, being found in 14 of the 41 clinical isolates across 6 of the 16 patients, indicating strong adaptation and persistence in the respiratory tracts of patients with CF. Although ST309 was the most prevalent, ST167 was also predominant (accounting for 8 out of 41 isolates); the remaining ST types were found in smaller proportions. Collectively, the predominance of these two clones accounts for more than half of the studied strains (22 out of 41), highlighting their adaptability in patients with CF (Figure 1a).

**Fig. 1.**
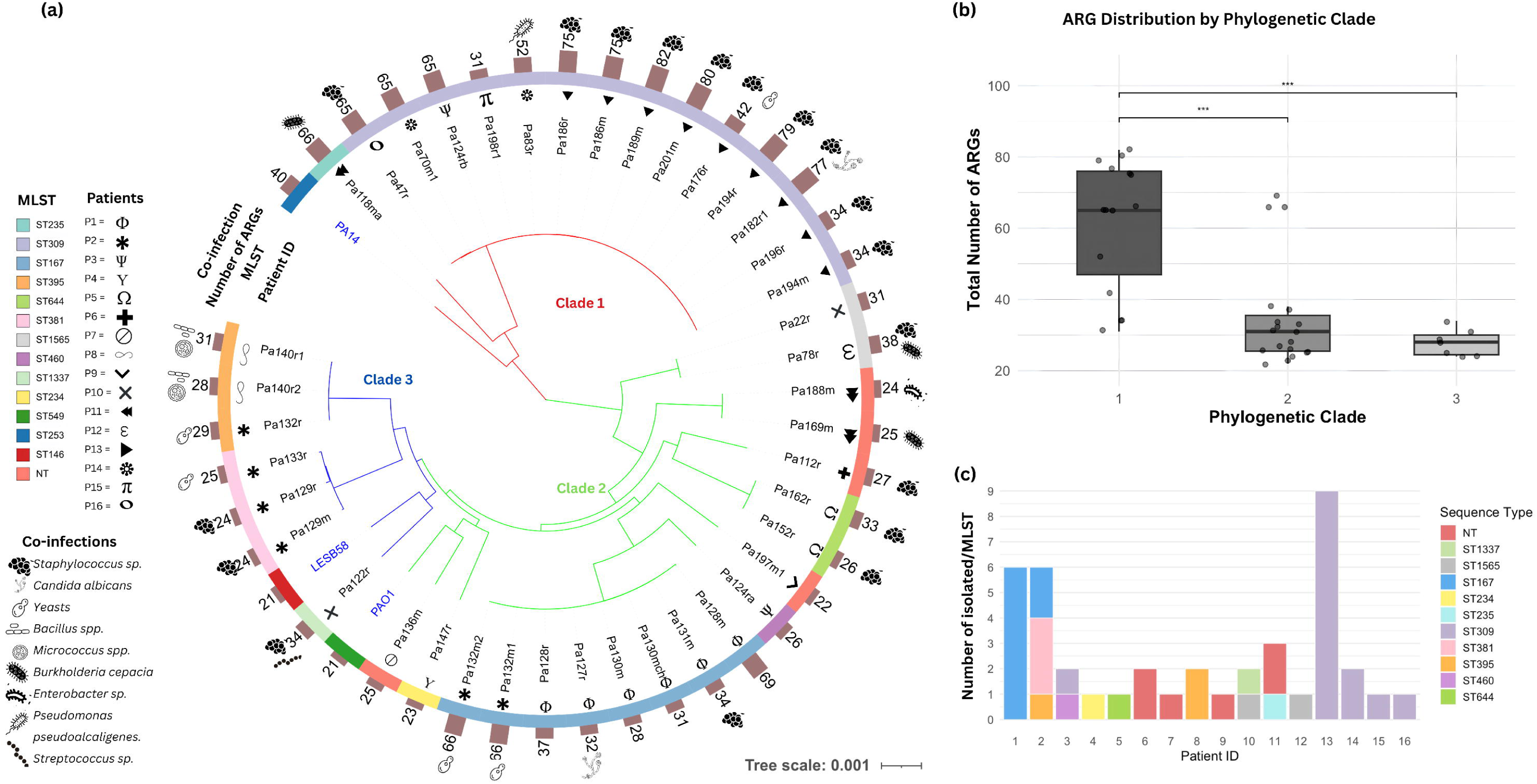
Phylogenetic analysis and resistome distribution of clinical *P. aeruginosa* isolates. (a) Maximum-likelihood phylogeny of the core genome of 41 P. aeruginosa isolates and the reference strains PAO1, PA14, and LESB58 (in blue). The symbols inside indicate the patient’s identification number (P1–P16), the colored bands represent the MLST types; the numbered bars represent the abundance of ARGs (antibiotic resistance genes), and the symbols on the surface represent the coinfection profile. (b) Box-plot comparison of the total number of ARGs identified across the three main phylogenetic clades; asterisks indicate statistically significant differences (*p* < 0.001). (c) Distribution of *P. aeruginosa* STs among the 16 patients.

The maximum-likelihood core genome phylogenetic tree shows that 41 *P. aeruginosa* strains group into three clades. Clade 1 was primarily formed by strains belonging to ST309, demonstrating the clonal expansion of this lineage within our cohort. Clade 2 exhibited greater genetic diversity, including most varieties of STs; meanwhile, Clade 3 consisted mainly of ST395 and ST381. Five genomes could not be assigned to existing STs (Pa169m, Pa188m, Pa136m, Pa112r, and Pa197m1), suggesting the presence of novel lineages in CF patients from Mexico. These isolates were obtained from four different patients and clustered within Clade 2 (Figure 1b, 1c).

### Lineage-specific distribution of resistome profiles

Antibiotic resistance determinants were identified using the CARD and NDARO databases via the BV-BRC platform, with results further validated by sraX. The analysis revealed significant heterogeneity in the resistome among the 41 *P. aeruginosa* clinical isolates. Detected genes were categorized by their mechanisms of resistance, including aminoglycoside-modifying enzymes, β-lactamases, efflux pump systems, target-site modifications (fluoroquinolones), and sulfonamide and phenicol resistance, among others (Figure 2a).

**Fig. 2.**
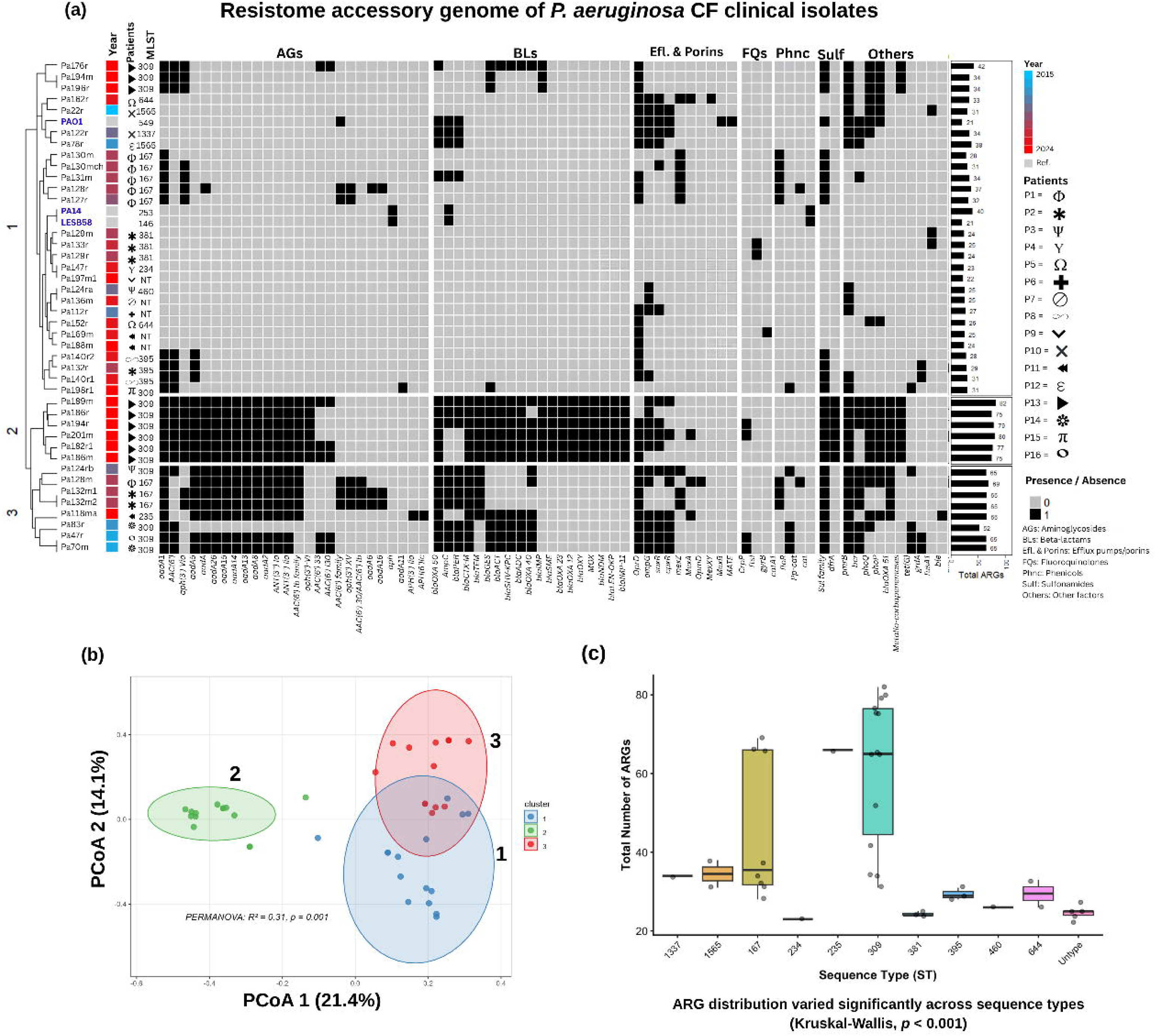
Resistome profile of clinical isolates of *P. aeruginosa*. (a) Binary heat map showing the presence (black) or absence (white) of antibiotic resistance genes (ARGs) in the 41 clinical isolates of *P. aeruginosa*. The isolates are sorted according to their resistome cluster (left axis). The heat map is stratified by class of antimicrobials: aminoglycosides (AG), β-lactams (BL), efflux pumps and porins (Efl. and Porins), fluoroquinolones (FQs), phenolics (Phnc), sulfonamides (Sulf), and other resistance factors. The colored columns represent the year of isolation; the symbols represent the patient to whom each isolate belongs; and the MLST is identified by letters. The bars on the right represent the richness of antibiotic resistance genes, including those comprising the core genome. (b) Principal coordinate analysis (PCoA) based on the Bray–Curtis dissimilarity of the resistome, showing three distinct clusters (PERMANOVA, R² = 0.31, p = 0.001). (c) Box plot showing the total number of ARGs identified in the different sequence types (STs); statistical significance was determined using the Kruskal–Wallis test (*p* < 0.001).

Resistome analysis identified 109 resistance genes in the 41 clinical isolates, of which 23 constituted the core resistome (present in at least 80% of the strains) (Table 2). Figure 2a shows a presence-absence heatmap of 86 antibiotic resistance genes (ARGs) representing the accessory resistome of the 41 clinical isolates, compared against the reference strains PAO1, PA14, and LESB58. To facilitate the analysis of resistance patterns, genes were grouped by their mechanism of action and protein family, including aminoglycoside-modifying enzymes, efflux systems, β-lactamases, fluoroquinolone-modifying enzymes, phenicols, sulfonamides, and other resistance determinants.

**Table 2.** Core resistome genes identified in. ≥ 80% of *Pseudomonas aeruginosa* isolated

| Gene category | Gene name | Prevalence % | Function |
| --- | --- | --- | --- |
| Fosfomycin | *fosA* | 95.5 | Modifying enzyme |
| Aminoglycosides | *aph(3')-Iib* | 95.5 | Modifying enzyme |
| Polymyxin | *arnA* | 97.7 | Modification of lipid A |
| β-lactams | *blaOXA* | 93.2 | Hydrolytic enzyme |
|  | *blaPDC* | 93.2 | Hydrolytic enzyme |
| Efflux Pumps/Porins | *mexB* | 100.0 | MexAB-OprM system |
|  | *mexJ* | 100.0 | MexCD-OprJ system |
|  | *mexH* | 97.7 | MexGHI-OpmD system |
|  | *mexM* | 97.7 | MexJK-MexM system |
|  | *mexP* | 100.0 | MexPQ-OpmE system |
|  | *mexD* | 100.0 | MexCD-OprJ system |
|  | *mexF* | 100.0 | MexEF-OprN system |
|  | *mexV* | 100.0 | MexVW-OprM system |
|  | *mexY* | 97.7 | MexXY-OprM system |
|  | *muxB* | 100.0 | MuxABC-OpmB system |
|  | *muxC* | 100.0 | MuxABC-OpmB system |
|  | *triAB* | 100.0 | TriABC-OpmH system |
| Fluoroquinolones | *parC* | 100.0 | Targets of quinolones. |
|  | *parE* | 100.0 | Targets of quinolones. |
|  | *gyrA* | 100.0 | Targets of quinolones. |
| Other determinants | *23SrRNA* | 97.7 | Components of the bacterial ribosome |
|  | *16SrRNA* | 97.7 |  |
| Phenicol | *catB7* | 90.9 | Modifying enzyme |

**Table 3.** Per-isolate genotype–phenotype concordance in *Pseudomonas aeruginosa* clinical isolates (n=41) , sorted by patient. First letter, phenotypic result; second letter, genomic prediction based on resistance gene presence/absence. R/R, concordant resistant; S/S, concordant susceptible; R/S, phenotypically resistant without detected gene; S/R, gene detected in phenotypically susceptible isolate. PEN, penicillins; CEP, cephalosporins; CAR, carbapenems; MON, monobactams; AGly, aminoglycosides; FLQ, fluoroquinolones; PLM, polymyxins.

| Genome | Patient | Year | AMR Profile | Phenotype / Genotype by drug class |  |  |  |  |  |  | Overall concordance |
| --- | --- | --- | --- | --- | --- | --- | --- | --- | --- | --- | --- |
|  |  |  |  | PE N | CEP | CA R | MON | AGI y | FLQ | PL M |  |
| *Pa127r* | PX1 | 2020 | MDR | S/R | S/R | S/R | S/S | R/R | R/R | R/S | **42.9%** |
| *Pa128m* |  | 2021 | Non-MDR | S/R | S/R | S/R | S/S | R/R | R/R | S/R | **42.9%** |
| *Pa128r* |  | 2021 | Non-MDR | S/R | S/R | S/R | S/S | R/R | R/R | S/S | **57.1%** |
| *Pa130m* |  | 2021 | MDR | S/R | S/R | R/S | S/S | R/R | R/R | S/S | **57.1%** |
| *Pa130mch* |  | 2021 | MDR | S/R | S/R | R/S | S/S | R/R | R/R | S/S | **57.1%** |
| *Pa131m* |  | 2021 | MDR | S/R | S/R | R/R | S/S | R/R | R/R | S/S | **71.4%** |
| *Pa129m* | PX2 | 2021 | MDR | R/R | R/R | R/S | S/S | R/R | R/R | S/S | **85.7%** |
| *Pa129r* |  | 2021 | XDR | R/R | R/R | R/S | R/S | R/R | R/R | S/S | **71.4%** |
| *Pa132m1* |  | 2021 | Non-MDR | S/R | S/R | S/R | S/S | R/R | R/R | S/R | **42.9%** |
| *Pa132m2* |  | 2021 | Non-MDR | S/R | S/R | S/R | S/S | R/R | R/R | S/R | **42.9%** |
| *Pa132r* |  | 2021 | MDR | R/R | R/R | R/R | R/S | R/R | S/R | S/S | **71.4%** |
| *Pa133r* |  | 2022 | XDR | R/R | R/R | R/S | R/S | R/R | R/R | S/S | **71.4%** |
| *Pa124ra* | PX3 | 2019 | XDR | R/R | R/R | R/S | R/S | R/R | R/R | S/R | **57.1%** |
| *Pa124rb* |  | 2019 | MDR | R/R | R/R | R/R | S/S | R/R | R/R | S/R | **85.7%** |
| *Pa147r* | PX4 | 2023 | MDR | R/R | R/R | R/S | S/S | R/R | S/R | S/S | **71.4%** |
| *Pa152r* | PX5 | 2023 | MDR | R/R | R/R | S/R | R/S | S/R | R/R | S/S | **57.1%** |
| *Pa162r* |  | 2023 | MDR | R/R | R/R | S/R | R/S | S/R | R/R | S/R | **42.9%** |
| *Pa112r* | PX6 | 2018 | MDR | R/R | R/R | R/R | R/S | R/R | S/R | S/R | **57.1%** |
| *Pa136m* | PX7 | 2023 | Non-MDR | S/R | S/R | S/S | S/S | S/R | S/R | S/R | **28.6%** |
| *Pa140r1* | PX8 | 2023 | MDR | R/R | R/R | R/R | S/S | R/R | R/R | S/R | **85.7%** |
| *Pa140r2* |  | 2023 | MDR | R/R | S/R | R/R | S/S | R/R | R/R | S/S | **85.7%** |
| *Pa197m1* | PX9 | 2024 | Non-MDR | S/R | S/R | S/S | S/S | S/R | S/R | S/S | **42.9%** |
| *Pa122r* | PX10 | 2019 | Non-MDR | R/R | S/R | S/R | S/S | R/R | S/R | S/R | **42.9%** |
| *Pa22r* |  | 2015 | MDR | R/R | S/R | R/R | S/S | S/R | R/R | S/R | **57.1%** |
| *Pa118ma* | PX11 | 2024 | MDR | S/R | R/R | S/R | S/S | R/R | R/R | S/R | **57.1%** |
| *Pa169m* |  | 2023 | Non-MDR | S/R | S/R | S/R | S/S | R/R | R/R | S/S | **57.1%** |
| *Pa188m* |  | 2024 | MDR | S/R | R/R | S/R | S/S | R/R | R/R | S/S | **71.4%** |
| *Pa78r* | PX12 | 2017 | Non-MDR | S/R | S/R | S/R | S/S | R/R | R/R | S/R | **42.9%** |
| *Pa176r* | PX13 | 2024 | XDR | R/R | R/R | R/R | R/R | R/R | R/R | S/R | **85.7%** |
| *Pa182r1* |  | 2024 | XDR | R/R | R/R | R/R | R/R | R/R | R/R | S/R | **85.7%** |
| *Pa186m* |  | 2024 | XDR | R/R | R/R | R/R | R/R | R/R | R/R | S/R | **85.7%** |
| *Pa186r* |  | 2024 | XDR | R/R | R/R | R/R | R/R | R/R | R/R | S/R | **85.7%** |
| *Pa189m* |  | 2024 | XDR | R/R | R/R | R/R | R/R | R/R | R/R | S/R | **85.7%** |
| *Pa194m* |  | 2024 | XDR | R/R | R/R | R/R | R/R | R/R | R/R | S/R | **85.7%** |
| *Pa194r* |  | 2024 | XDR | R/R | R/R | R/R | R/R | R/R | R/R | S/R | **85.7%** |
| *Pa196r* |  | 2024 | XDR | R/R | R/R | R/R | R/R | R/R | R/R | S/R | **85.7%** |
| *Pa201m* |  | 2024 | XDR | R/R | R/R | R/R | R/R | R/R | R/R | S/R | **85.7%** |
| *Pa70m1* | PX15 | 2016 | XDR | R/R | R/R | R/R | R/S | R/R | R/R | S/R | **71.4%** |
| *Pa83r* | PX14 | 2017 | XDR | R/R | R/R | R/R | R/S | R/R | R/R | S/R | **71.4%** |
| *Pa198r1* | PX15 | 2024 | XDR | R/R | R/R | R/S | R/S | R/R | R/R | S/R | **57.1%** |
| *Pa47r* | PX16 | 2016 | MDR | R/R | R/R | S/R | S/S | R/R | S/R | S/R | **57.1%** |
| **Concordance by drug class** |  |  |  | **65.9** | **63.4** | **48.8** | **75.6** | **87.8** | **82.9** | **34.1** | **Global:** |
|  |  |  |  | **%** | **%** | **%** | **%** | **%** | **%** | **%** | **65.5%** |

A principal coordinate analysis (PCoA) was performed based on the Hamming distances between the genotypic resistance profiles. This analysis confirmed the existence of three distinct genomic groups—hereafter referred to as resistome clusters to distinguish them from the phylogenetic clades—among the genomes analyzed here (*PERMANOVA*: R^2^= 0.31, *p* = 0.001).

Notably, these clusters revealed a non-random distribution of resistance determinants, with a significantly greater accumulation of antibiotic resistance genes (ARGs) observed in the genomes belonging to cluster 2, highlighting how specific gene families drive differentiation in resistance profiles (Figure 2b).

A strong association was observed between sequence types (STs) and the accumulation of ARGs (Figure 2c). The ST309 lineage exhibited the highest resistance burden, with a median of 66.0 ARGs (range: 31–82), significantly exceeding that of the other identified groups (*p* < 0.001). Other high-risk lineages, such as ST167 and ST1565, had medians of 35.5 and 34.5 ARGs, respectively, although with a resistance burden significantly lower than that observed in ST309. These results highlight a lineage-dependent distribution of resistance determinants, confirming ST309 as the primary reservoir of the multidrug-resistance profiles observed in our cohort (Table 4).

**Table 4.** Distribution and statistical significance of resistome profiles among different *P. aeruginosa* sequence types (STs).

| Lineage (ST) | Isolates (n) | Median ARG Count (IQR) | ARG Range | Statistical Significance (p-adj)* |
| --- | --- | --- | --- | --- |
| ST309 | 14 | 66.0 (65–79) | 31–82 | Reference |
| ST167 | 8 | 35.5 (28–66) | 28–69 | $p < 0.001$ |
| ST395 | 3 | 29.0 (28–31) | 28–31 | $p < 0.001$ |
| ST381 | 3 | 24.0 (24–25) | 24–25 | $p < 0.001$ |
| ST644 | 2 | 29.5 (26–33) | 26–33 | $p < 0.001$ |
| ST1565 | 2 | 34.5 (31–38) | 31–38 | $p < 0.001$ |
| Others | 4 | 26.5 (24–31) | 23–66 | $p < 0.001$ |
| Untype | 5 | 25.0 (24–25) | 22–27 | $p < 0.001$ |

\*Kruskal-Wallis global test $p < 0.001$ ; pairwise comparisons performed using Dunn's test with Holm-Bonferroni adjustment against ST309 as the reference lineage.

Similarly, single-nucleotide polymorphisms (SNPs) and insertion/deletion mutations were identified in several regulatory and structural genes. All 41 isolates analyzed harbored mutations in *gyrA* and *parE*, the genes encoding target sites for fluoroquinolones. Additionally, 17 isolates presented mutations in the porin gene *oprD*, typically associated with carbapenem resistance. To a lesser extent, mutations were identified in regulators of efflux pumps and two-component systems, such as *mexZ*, *nalD*, *phoP*, *phoQ*, and *nalC*. These mutational profiles likely contribute to the clinical persistence of *P. aeruginosa* in patients undergoing long-term fluoroquinolone therapy, further complicating therapeutic management.

To better understand the evolutionary factors that determine the observed resistance profiles, we conducted a comparative analysis stratified by phylogenetic clades. As shown in Table 5, the ARG burden showed significant stratification among groups (Kruskal-Wallis, *p* < 0.001), with Clade 1 exhibiting a notably higher resistance burden (mean: 75 ARGs/genome) compared to Clades 2 and 3 (means: 29.5 and 26.5, respectively; *p* < 0.001). These results not only reinforce the previously identified association between specific lineages (STs) and ARG accumulation but also identify Clade 1 lineages as the primary reservoir of multidrug resistance to antibiotics in our study population.

**Table 5.** Distribution and statistical significance of resistome profiles among three clades of P. aeruginosa.

| Clade | Isolates (n) | Median ARG Count (IQR) | ARG Range (Min–Max) | $p$-value* |
| --- | --- | --- | --- | --- |
| Clade 1 | 13 | 75.0 (66–80) | 31–82 | Reference |
| Clade 2 | 18 | 29.5 (25–33) | 23–69 | $p < 0.001$ |
| Clade 3 | 10 | 26.5 (25–31) | 24–34 | $p < 0.001$ |

\*The statistical significance was calculated using the Kruskal-Wallis test, followed by Dunn's post-hoc test, comparing each clade against Clade 1.

Analysis of resistance-associated mutations revealed variable frequencies among target genes in the 41 *P. aeruginosa* clinical isolates (Figure 3). Substitutions within the quinolone resistance-determining regions (QRDR) were ubiquitous, with both *gyrA* (n=41,100%) and *parE* (n=41,100%) harboring mutations across the entire cohort. High mutation frequencies were also identified in genes regulating carbapenem permeability and polymyxin adaptation, including *oprD* (n=17,41.5%), *phoQ* (n=17,41.5%), and *phoP* (n=15,36.6%). In contrast, mutations occurred at lower rates among multidrug efflux pump transcriptional regulators, specifically *nalC* (n=4, 9.8%), *nalD* (n=2, 4.9%), and *mexZ* (n=2, 4.9%) (Supplementary Table S2).

**Fig. 3.**
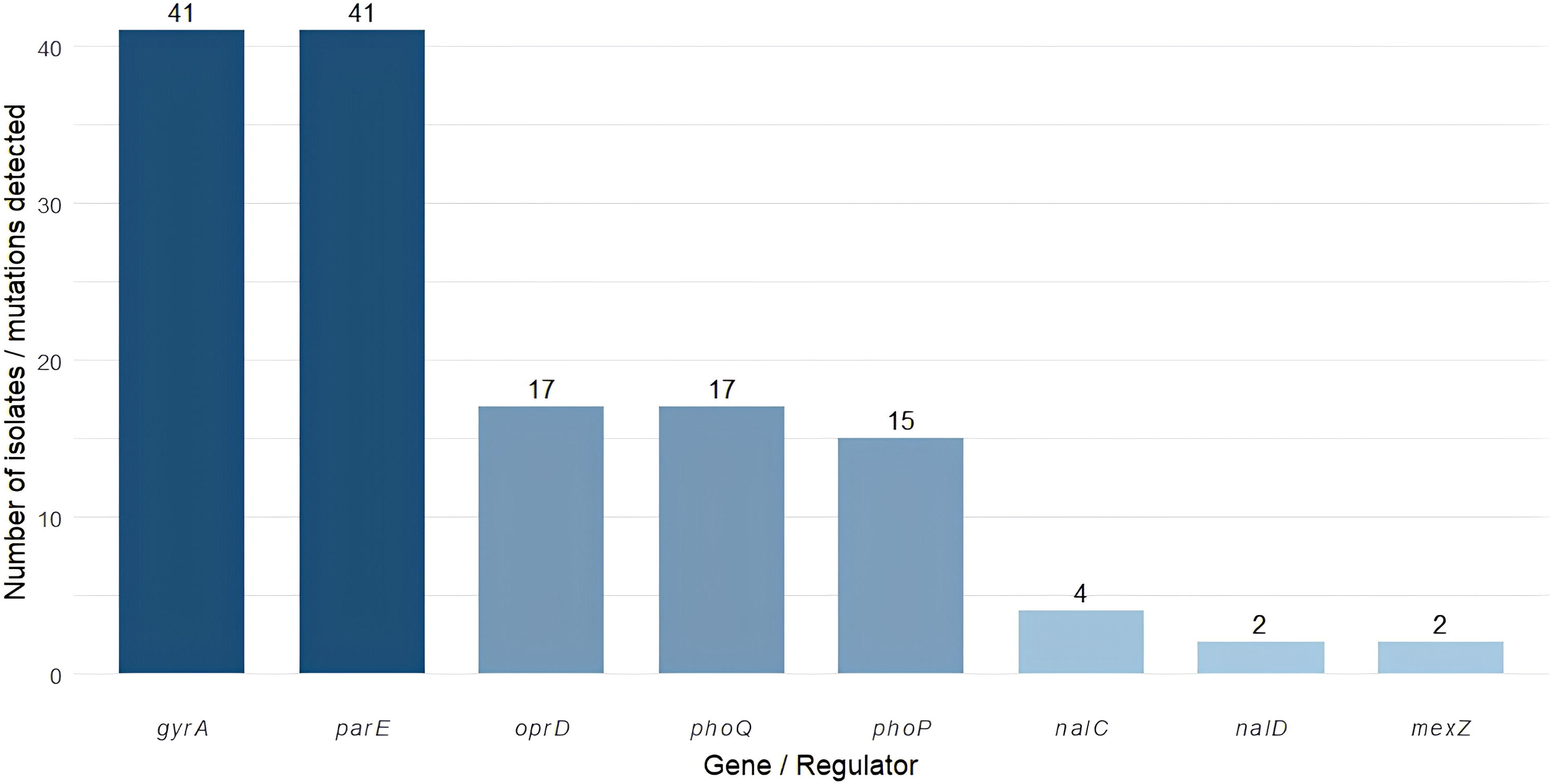
Prevalence of resistance-conferring mutations in *P. aeruginosa* clinical isolates. Bar chart showing the frequency of detected mutations in key resistance-associated genes and regulatory loci. The y-axis represents the number of isolates (n=41) harboring mutations in the specified genes (x-axis). Darker bars indicate high-frequency mutations present in all analyzed isolates (n=41), while lighter shades represent decreasing mutation prevalence across the clinical cohort.

### Genotype–phenotype concordance

Genotype–phenotype concordance was assessed by comparing susceptibility results with genomic resistance determinants identified by sraX analysis across 41 clinical *P. aeruginosa* isolates (Table 3). Concordance was defined as phenotypic resistance accompanied by at least one corresponding resistance gene, or phenotypic susceptibility in the absence of such genes. Intrinsic determinants (*ampC*, *blaADC*) and multi-drug efflux pump genes (*mexAB-oprM, mexXY-oprM, mexCD-oprJ, mexEF-oprN*) were excluded, as they preclude class-specific attribution in the absence of expression data. Global concordance across all seven drug classes was 65.5%.

Aminoglycosides and fluoroquinolones showed the highest concordance (87.8% and 82.9%, respectively), with no R/S discordances detected in either class. Aminoglycoside resistance was supported by a broad repertoire of aminoglycoside-modifying enzyme (AME) genes, including acetyltransferases (*aac(6*′*)-Ib, aac(6*′*)-Ib-cr, aac(3)-Ia*), nucleotidyltransferases (*ant(3*″*)-Ia, ant(3*″*)-IIa*), phosphotransferases (*aph(3*′*)-Iia, aph(3*′*)-Iib, aph(6)-Ic*), and *aadA* variants. Fluoroquinolone resistance was associated with mutations in *gyrA, gyrB, parC*, and *parE*, together with the plasmid-mediated enzyme *crpP* and the efflux regulator *soxR*. The S/R discordances observed in both classes (5 and 7 isolates, respectively) likely reflect subthreshold gene expression or the requirement for co-occurring regulatory mutations.

Monobactam (aztreonam) concordance was 75.6%. Given the absence of class-specific acquired determinants for aztreonam in *P. aeruginosa*, *blaIMP* and *blaNDM* were included on the basis of their documented hydrolytic activity against this compound. Nine of 19 phenotypically resistant isolates carried these genes (R/R), while the remaining ten R/S discordances most plausibly reflect *MexAB-OprM* or *MexXY-OprM* overexpression — efflux mechanisms excluded from this analysis due to their multi-drug substrate profile.

Penicillin and cephalosporin concordance were 65.9% and 63.4%, respectively, with all discordances exclusively of the S/R type (14 and 15 isolates, respectively) and no R/S cases in either class. The primary determinant identified was *blaPDC*, an inducible chromosomal AmpC-type β-lactamase ubiquitous in *P. aeruginosa,* accompanied by acquired enzymes including *blaGES, blaPER, blaTEM, blaCTX-M, blaACT, MOX*, and *blaOXY*. The predominance of S/R discordances is consistent with the inducible nature of *blaPDC*, whereby constitutive high-level expression requires loss-of-function mutations in *ampD* or *dacB* that cannot be inferred from gene presence alone.

Carbapenem concordance was the lowest among β-lactam classes (48.8%), with discordances in both directions. The 18 R/R isolates harboured *blaIMP*, *blaIMP-11, blaNDM*, and/or *oprD* variants, while the eight R/S cases — resistant without a detectable determinant — likely reflect OprD downregulation or frameshift mutations beyond the reference dataset, potentially compounded by *MexAB-OprM* upregulation. The 13 S/R cases suggest that carbapenemase gene carriage alone is insufficient to predict resistance, as expression levels, enzyme kinetics, and compensatory permeability changes collectively determine the phenotypic outcome — consistent with the widely reported complexity of carbapenem resistance prediction in *P. aeruginosa*.

Polymyxins showed the lowest concordance overall (34.1%), with 26 of 41 isolates exhibiting S/R discordance. Only *pmrB* was included as a specific acquired determinant, as the regulatory genes *arnA, phoP*, and *phoQ* require gain-of-function mutations — not mere presence — to activate lipid A modification and confer resistance. The high S/R rate reflects the core-genome nature of *pmrB* in *P. aeruginosa*, underscoring the need for SNP-level analysis of two-component system regulators rather than presence/absence screening for accurate polymyxin resistance prediction.

At the per-isolate level, concordance ranged from 28.6% to 85.7%. Patient PX13 (n=9 isolates, all XDR, 2024) showed uniformly high concordance (85.7%), with resistance genotypically confirmed across six of seven drug classes in every isolate and S/R discordance restricted to polymyxins, suggesting clonal persistence of a single XDR lineage. Patient PX2 (n=6 isolates, Non-MDR to XDR) showed the greatest intra-patient heterogeneity: R/S discordances in carbapenems were detected exclusively in the XDR isolates (Pa129r, Pa133r) and not in the MDR or Non-MDR isolates from the same patient, indicating acquisition of non-enzymatic — likely OprD-mediated — carbapenem resistance during clinical evolution under selective pressure; additionally, Pa132r from this same patient exhibited R/S discordance in monobactams and S/R in fluoroquinolones, further reflecting the divergent resistance trajectories within a single host. Patient PX8 (n=2 isolates, both MDR, 2023) achieved the second highest concordance (85.7%), consistent with a well-established MDR profile genotypically supported across all major drug classes. The lowest per-isolate concordance was observed in Pa136m (PX7, 28.6%), which carried resistance genes for five of seven drug classes yet remained phenotypically Non-MDR across all, a genotypically loaded but phenotypically quiescent profile potentially attributable to regulatory silencing warranting further transcriptomic investigation.

### Antimicrobial susceptibility profiles and resistance patterns

Phenotypic antimicrobial susceptibility profiles of the 41 *P. aeruginosa* clinical isolates are shown in Figure 4. Hierarchical clustering based on resistance profile similarity (Ward linkage, Hamming distance) resolved the collection into two phenotypically distinct populations. The upper cluster comprised isolates with predominantly susceptible profiles, with resistance largely restricted to aminoglycosides and fluoroquinolones, while the lower cluster was characterized by extensive resistance spanning multiple drug classes, including β-lactams, aminoglycosides, and fluoroquinolones. This bimodal distribution suggests the co-circulation of two epidemiologically distinct *P. aeruginosa* populations within the clinical setting.

**Fig. 4.**
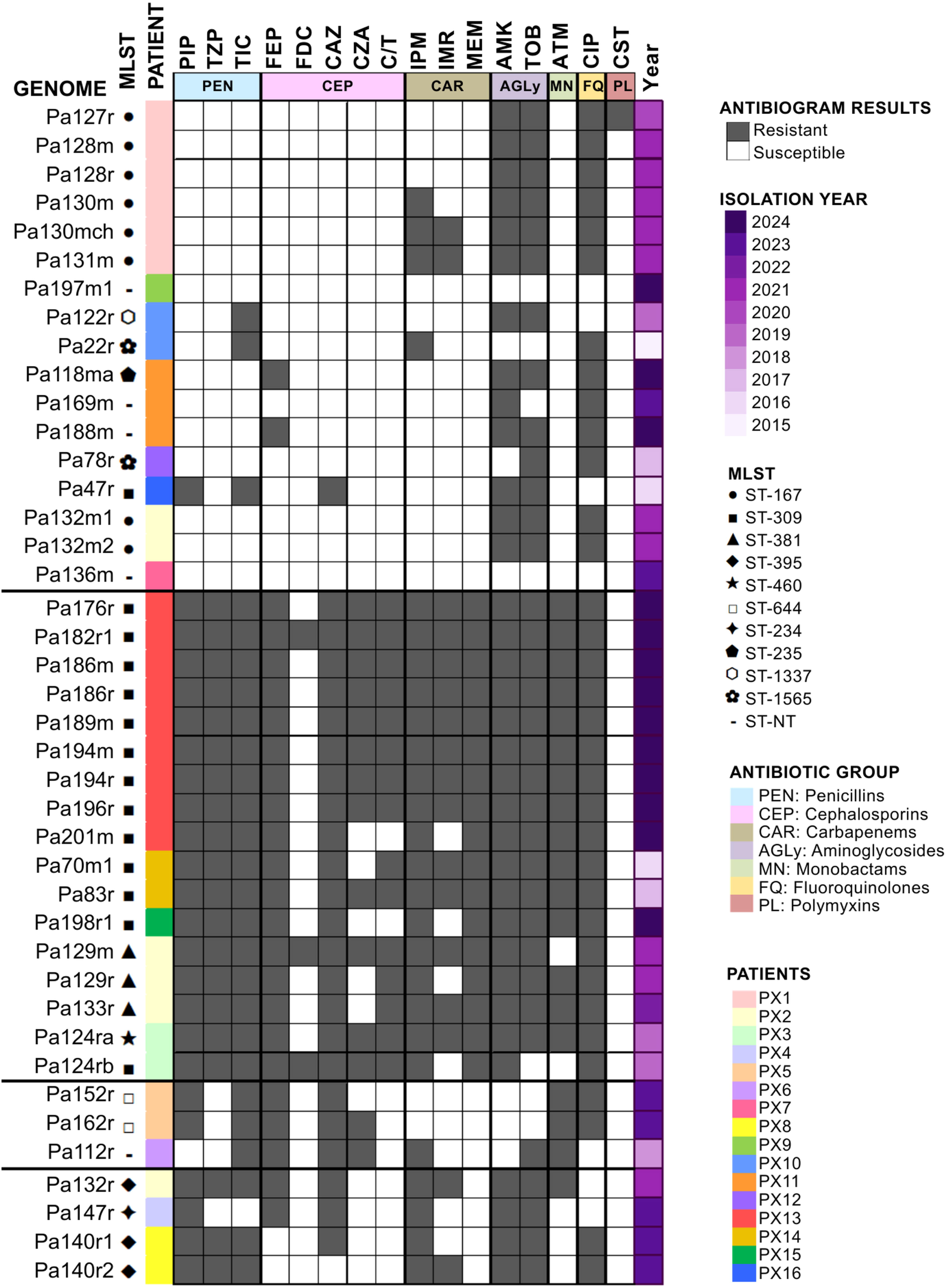
Antimicrobial susceptibility profiles of *P. aeruginosa* clinical isolates (n=41). Black cells indicate phenotypic resistance; grey cells indicate susceptibility. Isolates are grouped by hierarchical clustering (Ward linkage, Hamming distance) based on resistance profile similarity. The MLST symbol denotes the sequence type of each isolate. The isolation year is represented by a purple gradient (darkest, 2024; lightest, 2015). Row colours indicate the patient of origin. Antibiotic abbreviations: PIP, piperacillin; TZP, piperacillin-tazobactam; TIC, ticarcillin-clavulanate; FEP, cefepime; FDC, cefiderocol; CAZ, ceftazidime; CZA, ceftazidime-avibactam; C/T, ceftolozane-tazobactam; IPM, imipenem; IMR, imipenem-relebactam; MEM, meropenem; AMK, amikacin; TOB, tobramycin; ATM, aztreonam; CIP, ciprofloxacin; CST, colistin. PEN, penicillins; CEP, cephalosporins; CAR, carbapenems; AGly, aminoglycosides; MN, monobactams; FQ, fluoroquinolones; PL, polymyxins.

**Fig. 5.**
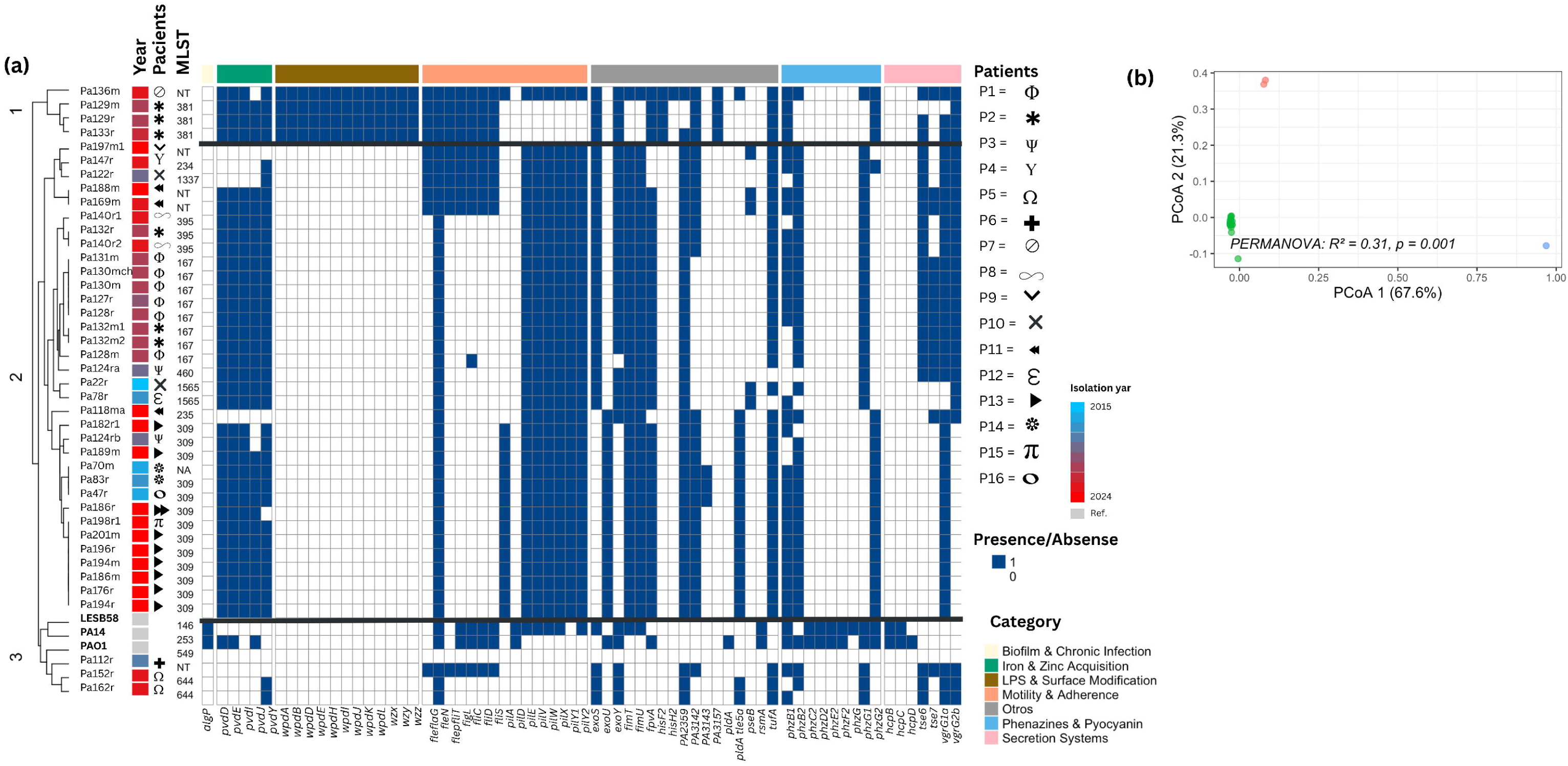
Virulome profiling of *P. aeruginosa* clinical isolates. (a) Heat map illustrating the presence (blue) or absence (white) of virulence-associated genes across 41 *P. aeruginosa* clinical isolates, clustering based on their virulome profiles. Genes are color-coded by functional category: biofilm and chronic infection, iron and zinc acquisition, lipopolysaccharide (LPS) and surface modification, motility and adherence, phenazines and pyocyanin, secretion systems, and other virulence factors. Sidebars indicate the isolation year and the corresponding sequence type (ST) and patient ID for each isolate. (b) Principal Coordinates Analysis (PCoA) based on Bray– Curtis dissimilarity of the virulome profiles, showing distinct clustering patterns among isolates (PERMANOVA, 0.31, 0.001).

Among individual antibiotic agents, cefiderocol retained susceptibility in the largest proportion of isolates, including several classified as XDR, positioning it as one of the few therapeutic options with preserved activity across the collection. In contrast, colistin resistance was detected in a subset of isolates from the upper cluster that remained susceptible to most other drug classes, indicating that polymyxin resistance can emerge independently of a broader MDR or XDR phenotype in this collection. Comparison of imipenem and imipenem-relebactam profiles revealed that the addition of relebactam restored susceptibility in a subset of imipenem-resistant isolates, consistent with the documented activity of this combination against *OprD*-deficient or *MexAB-OprM*-overexpressing strains in the absence of acquired carbapenemases.

Aminoglycosides and fluoroquinolones showed the most homogeneous within-cluster distribution, with near-complete resistance in the lower cluster and predominant susceptibility in the upper cluster, reflecting the high genotype–phenotype concordance observed for these drug classes (87.8% and 82.9%, respectively; Table 3).

A temporal trend was evident in the resistance profiles: isolates collected in 2023–2024 were disproportionately represented in the lower, high-resistance cluster, whereas isolates from 2015– 2018 were more evenly distributed across both clusters. This pattern suggests a progressive shift towards more resistant phenotypes over the surveillance period, consistent with the accumulation of resistance determinants under sustained antibiotic selective pressure.

At the patient level, two patterns of clinical relevance were identified. Patient PX13 contributed nine isolates, all collected in 2024 and all classified as XDR, exhibiting a completely homogeneous resistance profile across all nine isolates with resistance confirmed across six of seven drug classes. The absence of any phenotypic variation across these isolates strongly suggests clonal persistence of a single XDR lineage throughout the sampling period, a finding of significant clinical and infection control concern. Patient PX2, in contrast, contributed isolates distributed across both clusters, with Non-MDR and MDR isolates co-occurring alongside XDR isolates from the same patient. This intra-patient phenotypic divergence indicates the acquisition or emergence of additional resistance mechanisms during the clinical course, consistent with *P. aeruginosa* adaptive evolution under antibiotic pressure, and underscores the importance of sequential sampling in patients with recurrent or chronic *P. aeruginosa* infections.

### Virulome

Analysis of virulome profiles identified 67 distinct patterns, primarily categorized into motility and adhesion, lipopolysaccharide (LPS), and surface modification factors. Clade 1 was characterized by exclusive LPS and surface modification genes, whereas Clade 2 encompassed the majority of motility and adhesion-related genes. Notably, the *pilD* was found exclusively in reference genomes (LESB58 and PA14), while *fleN* was conserved across all clinical isolates. Interestingly, no clinical genomes contained biofilm-related genes, which were only present in the reference strains LESB58 and PA14, suggesting adaptive mechanisms in patients with CF, such as post-translational modifications or other mechanisms that lead to chronic persistence in the lung. Conversely, genes associated with iron and zinc acquisition were predominant in Clades 1 and 2. Distribution of phenazine and pyocyanin genes varied across the three clades; specifically, *phzB1*, *phzB2*, *phzC2*, *phzD2*, and *phzE2* were restricted to reference genomes. Finally, secretion systems showed heterogeneous distribution, with *hcpB*, *hcpC,* and *hcpD* identified exclusively in clinical isolates (Supplementary Table S3).

A PCoA analysis of the virulence profile was performed, confirming the presence of three major clades, which correlate with the resistome clades (*PERMANOVA*: R^2^= 0.31, *p* = 0.001).

These provide evidence that the identified clades have evolved specific adaptations in parallel with their resistome, posing a significant challenge for the clinical management of patients colonized by these high-risk sequence types.

### Mobilome analyses

The mobilome analysis demonstrated interactions among mobile genetic elements (MGEs), insertion sequences (ISs), and resistance genes in clinical isolates of *P. aeruginosa*. At the chromosomal level (n = 77), integrons and integrative conjugative elements (ICEs) were associated with resistance determinants for aminoglycosides (*aad*, *aph*, and *aac*), β-lactams, and phenicols. Plasmids (n = 18) and elements of uncertain localization (n = 1) were found to carry resistance genes, such as metallo-β-lactamases and aminoglycoside-modifying enzymes, flanked primarily by insertion sequences of the *IS6100* and *IS1* families. These results highlight the importance of resistance genes acquired through horizontal gene transfer, which modulate and expand the resistome of our cohort (Figure 6).

**Fig. 6.**
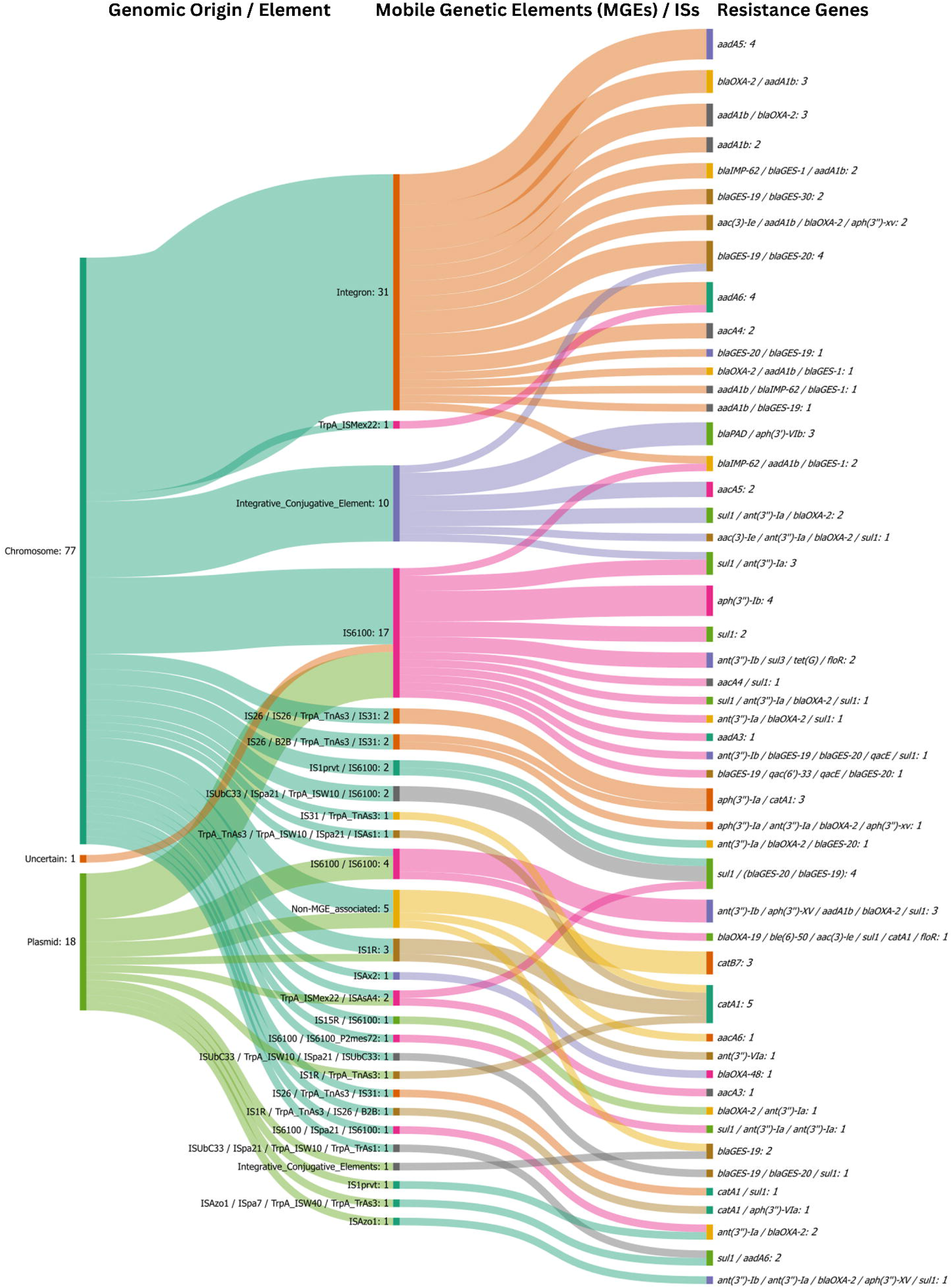
Sankey diagram illustrating the flow of mobile genetic elements and antimicrobial resistance genes in clinical isolates of *Pseudomonas aeruginosa*. MGEs associated with the chromosome (n = 77), plasmids (n = 18), and elements of uncertain localization (n = 1). The columns show, from left to right: level 1, the genomic element or origin; level 2, mobile genetic elements (MGEs/IS); and level 3, resistance genes. The thickness of the connections is proportional to the frequency of the observed associations.

## Discussion

The genomic analysis of *P. aeruginosa* isolates from paediatric CF patients in Mexico reveals a population structure dominated by a small number of highly adapted lineages, most notably ST309 and ST167. The predominance of ST309 across multiple patients and sampling years underscores its capacity for long-term persistence in CF airways, a pattern consistent with high-risk clones reported in other regions. Our phylogenomic reconstruction shows that ST309 isolates form a well-defined clade with limited intra-lineage diversity, suggesting either repeated acquisition of closely related strains or sustained within-host maintenance of a successful genotype. The identification of five untypeable isolates further highlights the presence of previously undescribed lineages circulating in this clinical setting, expanding the known genomic diversity of *P. aeruginosa* in Latin America. It is worth noting that ST309 was first reported in Mexico in 2017 as a high-risk clone in pediatric patients with bacteraemia, which underscores its significance due to its ability to cause hospital-acquired outbreaks and its capacity for adaptation and persistence in vulnerable populations [23]. Although other high-risk clones associated with nosocomial outbreaks have been reported in Mexico—such as ST244, ST235, and ST111— ST309 and ST167 predominated in our pediatric CF cohort [24].

Resistome profiling revealed substantial heterogeneity across isolates, yet also identified a conserved core of resistance determinants characteristic of *P. aeruginosa*. The exceptionally high ARG burden in ST309—more than double that of most other lineages—indicates strong lineage-specific enrichment of resistance determinants. This pattern aligns with reports describing ST309 as a globally emerging multidrug-resistant clone. The clear stratification of ARG counts across phylogenetic clades suggests that resistance accumulation is not random but shaped by lineage-specific evolutionary trajectories. The presence of rare determinants in only a few isolates, combined with the detection of novel allelic combinations, points to ongoing microevolution and sporadic acquisition events within the CF airway environment. The genomic diversity of resistance observed in our cohort is consistent with findings by Bianconi et al. [25], who demonstrated in a longitudinal CF study how persistent *P. aeruginosa* populations progressively accumulate resistance mutations, leading to MDR phenotypes and microevolutionary diversification within clonal lineages (ST390).

Mutation analysis further supports the notion of strong selective pressure imposed by chronic antimicrobial exposure. Universal mutations in *gyrA* and *parE* across all isolates indicate widespread fluoroquinolone adaptation, while disruptions in *oprD* and modifications in the *phoP/phoQ* two-component system reflect parallel evolution toward carbapenem and polymyxin tolerance. The relatively lower frequency of mutations in efflux regulators such as *mexZ*, *nalC*, and *nalD* suggests that regulatory rewiring is less common than target-site modification in this cohort, although their presence in a subset of isolates may contribute to multidrug resistance phenotypes. Similarly, Mattingly et al. (2025) [26] reported adaptive mutations in *phoPQ* among colistin-resistant *P. aeruginosa* isolates from CF patients, highlighting the strong evolutionary capacity of persistent clinical strains under polymyxin selective pressure. Regarding carbapenem resistance, a substantial proportion of isolates (41.5%) harbored alterations in the *oprD* porin gene [27]. Structural disruption or loss of function in OprD constitutes a well-established mechanism for imipenem and meropenem resistance in *P. aeruginosa* (Köhler et al., 1999), demonstrating that carbapenem resistance in our pediatric CF collection was primarily driven by chromosomal porin loss rather than acquired carbapenemases.

The genotype–phenotype concordance analysis highlights both the strengths and limitations of genomic prediction in *P. aeruginosa*. The 65.4% overall concordance reflects high accuracy for aminoglycosides and fluoroquinolones, where prediction is driven by established ARGs and target-site mutations. In contrast, the lower concordance observed for carbapenems (47.5%) and polymyxins (35.0%) underscores that resistance to these drug classes is largely driven by complex regulatory mutations—such as the constitutive induction of ampC/blaPDC, overactivation of the two-component system (e.g., phoPQ/pmrAB), or the loss of the oprD porin—rather than by the simple acquisition of transferable genes. This is best illustrated by isolate Pa136m, which displayed a high accumulation of ARGs across five drug classes yet remained phenotypically non-multidrug-resistant (Non-MDR). Such profiles, where genomic potential does not correlate with phenotypic susceptibility, suggest transcriptional silencing or loss-of-function regulatory mutations. These findings reinforce the need for integrated genomic and phenotypic surveillance, indicating that reliance on ARG presence/absence matrices alone may lead to an overestimation of resistance, necessitating the integration of SNP-level analysis and functional transcriptomics [29].

The virulome of the clinical isolates analyzed here showed the absence of biofilm-associated canonical genes, and the restriction of phenazine genes (*phzB1/B2, phzC2, phzD2, phzE2*) to the reference strains (PAO1 and PA14) reflects the counter-selection of acute virulence determinants during long-term persistence in the respiratory tract. The chronic adaptation of *P. aeruginosa* strains is characterized by the attenuation of metabolic and proinflammatory virulence factors to evade the host’s immune surveillance. Furthermore, we found structural components of the T6SS—*hcpB, hcpC, hcpD*—along with conserved iron and zinc acquisition pathways in all clinical clades, underscoring an evolutionary trade-off that prioritizes interbacterial competition and nutrient uptake over acute tissue damage within the polymicrobial airways of cystic fibrosis.

The virulome profiles observed in our cohort are consistent with the pathoadaptation of *P. aeruginosa* in the airways of patients with CF, as described by Winstanley, O’Brien, and Brockhurst (2016) [28]. During persistent infections, *P. aeruginosa* undergoes strong evolutionary trade-offs, modifying its virulence repertoire by attenuating highly immunogenic factors to evade host recognition. Simultaneously, the pathogen adapts to this environment through metabolic restructuring, hypermutability, and the transition to biofilm-associated phenotypes. Notably, while acute pathogenesis is often silenced, clinical strains retain mechanisms crucial for interbacterial competition, such as the T6SS, which ensures their ecological dominance and long-term persistence within the polymicrobial pulmonary niche.

Mobilome analysis demonstrates the mechanisms of resistance dissemination in clinical strains of *P. aeruginosa* in the context of cystic fibrosis, as high connectivity was observed between MGEs and ARGs. At the chromosomal level, we observed the recurrent presence of integrons and ICEs, indicating that a niche for the acquisition and integration of antimicrobial resistance cassettes exists in the lungs. Furthermore, the identification of plasmids carrying metallo-β-lactamases (*blaIMP*, *blaNDM*) associated with insertion sequences, such as IS6100, demonstrates the spread of carbapenem resistance determinants. These findings are consistent with evidence from other clinical settings with high selective pressure, where MGEs contribute to the transition toward multidrug-resistant (MDR) and extensively drug-resistant (XDR) phenotypes, complicating the therapeutic management of chronic infections.

Collectively, our results provide a detailed genomic portrait of *P. aeruginosa* circulating in Mexican paediatric CF patients, revealing dominant high-risk clones, lineage-specific resistance burdens, and recurrent adaptive mutations. This study provides essential baseline data for regional genomic surveillance and underscores the importance of monitoring both clonal dynamics and the evolution of resistance in chronic CF infections. The identification of novel lineages and diverse resistome profiles highlights the need for continued sequencing efforts to better understand the epidemiology and evolutionary potential of *P. aeruginosa* in Latin

America. Future work integrating transcriptomics, longitudinal sampling, and host-microbiome interactions will be critical to elucidate the mechanisms driving persistence and treatment failure in this clinically challenging pathogen.

## Conclusions

This study provides a high-resolution genomic portrait of *P. aeruginosa* circulating in paediatric CF patients in Mexico, revealing that long-term persistence in the CF airway is strongly associated with high-risk lineages, extensive antimicrobial resistance repertoires, and lineage-specific adaptive evolution. Among the isolates analyzed, ST309 and ST167 emerged as the dominant clones, each characterized by a substantial burden of resistance determinants and a marked ability to persist across multiple sampling years. Our findings demonstrate that the resistome of these lineages is not merely the result of sporadic gene acquisition, but reflects a dual architecture composed of a conserved core and a highly dynamic accessory genome that expands under chronic therapeutic pressure. These observations underscore the importance of implementing routine genomic surveillance to monitor clonal expansion and resistance evolution in paediatric clinical settings.

We further show that *P. aeruginosa* undergoes convergent pathoadaptation in response to sustained fluoroquinolone exposure, facilitating the transition from acute infection to chronic colonization. This evolutionary shift is evidenced by the attenuation of acute virulence traits alongside the retention of competitive functions such as the Type VI Secretion System. The widespread presence of chromosomal mutations in *oprD*, *phoP/phoQ*, and the quinolone resistance-determining regions of *gyrA* and *parE*, together with observed genotype–phenotype discordances, highlights that genomic predictions alone cannot fully capture the complexity of antimicrobial resistance in chronic CF infections. Instead, our results confirm that regulatory rewiring and structural gene modifications are the principal drivers of multidrug resistance in this context—mechanisms that frequently escape detection when relying solely on traditional ARG-based screening.

Overall, this work emphasizes the need for a multidisciplinary approach to managing chronic *P. aeruginosa* infections in paediatric CF patients. The persistence of extensively drug-resistant lineages such as ST309, combined with the emergence of resistance mechanisms beyond classical enzymatic pathways, calls for the integration of genomic surveillance, functional diagnostics, and the exploration of alternative therapeutic strategies. Strengthening these efforts will be essential to improving clinical outcomes and mitigating the long-term impact of multidrug-resistant *P. aeruginosa* on vulnerable paediatric populations.

## Author contributions

-Evelin Martínez-Rosales: Methodology, Investigation, Formal analysis, Data curation, Visualization, Writing—original draft.

-Rafael Coria Jiménez: Conceptualization, Funding acquisition, Supervision, Validation, Writing—review & editing.

-Armando Gerónimo-Gallegos: Investigation, Resources.

-Francisco Cuevas-Schacht: Investigation, Resources.

-Miriam Sarahi Lozano Gamboa: Methodology, Data curation, Formal analysis, Visualization.

-Marisol López-López: Supervision, Writing—review & editing.

-Rodolfo García-Contreras: Supervision, Writing—review & editing.

-Corina Diana Ceapă: Conceptualization, Methodology, Data curation, Project administration, Funding acquisition, Supervision, Validation, Writing—review & editing.

## Funding information

This work was supported by collaborations with MGI and Illumina for strain sequencing at no cost to the research group. We acknowledge receiving funding from the University of Illinois System and the Coordinación de la Investigación Científica of UNAM Seed Funding 2024 for the project “Targeting of mobile elements from multidrug-resistant ESKAPEE pathogens”. Additionally, this research was supported by the project INP-2026/011, which is funded by federal resources allocated to the Instituto Nacional de Pediatría. The funders had no role in study design, data collection and analysis, decision to publish, or preparation of the manuscript. *Ethical approval*

Respiratory samples were collected as part of routine clinical care and processed under institutional approval (INP-2026/011), following national regulations (NOM-012-SSA3-2012) and international guidelines for genomic data handling.

## Consent for publication

All authors have read and approved the final version of the manuscript and have given their explicit consent for its publication.

## Supporting information

Tablas complementarios

## Acknowledgements

The authors would like to thank the staff of the Experimental Bacteriology Laboratory of the Instituto Nacional de Pediatría, Mexico, for their support in the collection and processing of clinical isolates. This work was supported by the Doctoral fellowship (grant number 1329607) provided to E.M. by the Secretaría de Ciencia, Humanidades, Tecnología e Innovación (SECIHTI).

